# Design of the SARS-CoV-2 RBD vaccine antigen improves neutralizing antibody response

**DOI:** 10.1101/2021.05.09.443238

**Authors:** Thayne H. Dickey, Wai Kwan Tang, Brandi Butler, Tarik Ouahes, Sachy Orr-Gonzalez, Nichole D. Salinas, Lynn E. Lambert, Niraj H. Tolia

## Abstract

The receptor binding domain (RBD) of the SARS-CoV-2 spike protein is the primary target of neutralizing antibodies and is a component of almost all vaccine candidates. Here, RBD immunogens were created with stabilizing amino acid changes that improve the neutralizing antibody response, as well as characteristics for production, storage, and distribution. A computational design and *in vitro* screening platform identified three improved immunogens, each with approximately nine amino acid changes relative to the native RBD sequence and four key changes conserved between immunogens. The changes are adaptable to all vaccine platforms, are compatible with established changes in SARS-CoV-2 vaccines, and are compatible with mutations in emerging variants of concern. The immunogens elicit higher levels of neutralizing antibodies than native RBD, focus the immune response to structured neutralizing epitopes, and have increased production yields and thermostability. Incorporating these variant-independent amino acid changes in next-generation vaccines may enhance the neutralizing antibody response and lead to pan-SARS-CoV-2 protection.

## Main text

COVID-19 has emerged as one of the most significant global health issues and several SARS-CoV-2 vaccines have been successful in phase 3 trials and real-world deployment. Many of these vaccines contain an engineered SARS-CoV-2 spike protein antigen with stabilizing amino acid changes in the S2 domain^1–4^ that improve the efficacy^5–7^. However, next generation vaccines are necessary to address emerging strains, confer long-lasting protection, and overcome global distribution challenges. Stabilizing mutations in the spike protein outside of the S2 domain could further improve vaccine efficacy and stability to address these challenges.

The receptor binding domain (RBD) of the spike protein engages the Ace2 receptor to mediate viral entry, and it is consequently the target of the most potent SARS-CoV-2 neutralizing antibodies (Fig. 1a and b)^8–12^. In fact, the RBD alone is a vaccine candidate that elicits potent neutralizing antibodies and protective immunity in humans and animals^1,13–15^. In contrast, potent neutralizing antibodies are rare in convalescent patients relative to non-neutralizing antibodies that bind epitopes elsewhere on the spike protein^10–12^. This phenomenon could be due to structural instability in neutralizing epitopes of the RBD and the ability of the RBD to adopt a “down” conformation that obscures neutralizing epitopes in the spike trimer (Fig. 1a). The RBD is thought to transiently sample an “up” conformation compatible with Ace2 binding, which exposes neutralizing epitopes including some that may cross-protect against other coronaviruses^16–20^. Stabilizing the RBD structure, focusing the immune response to neutralizing epitopes and away from non-neutralizing epitopes, and exposing the RBD either as a stand-alone vaccine or in the RBD-up conformation of the spike trimer would be expected to increase the efficacy of spike-based vaccines.

**Fig. 1.**
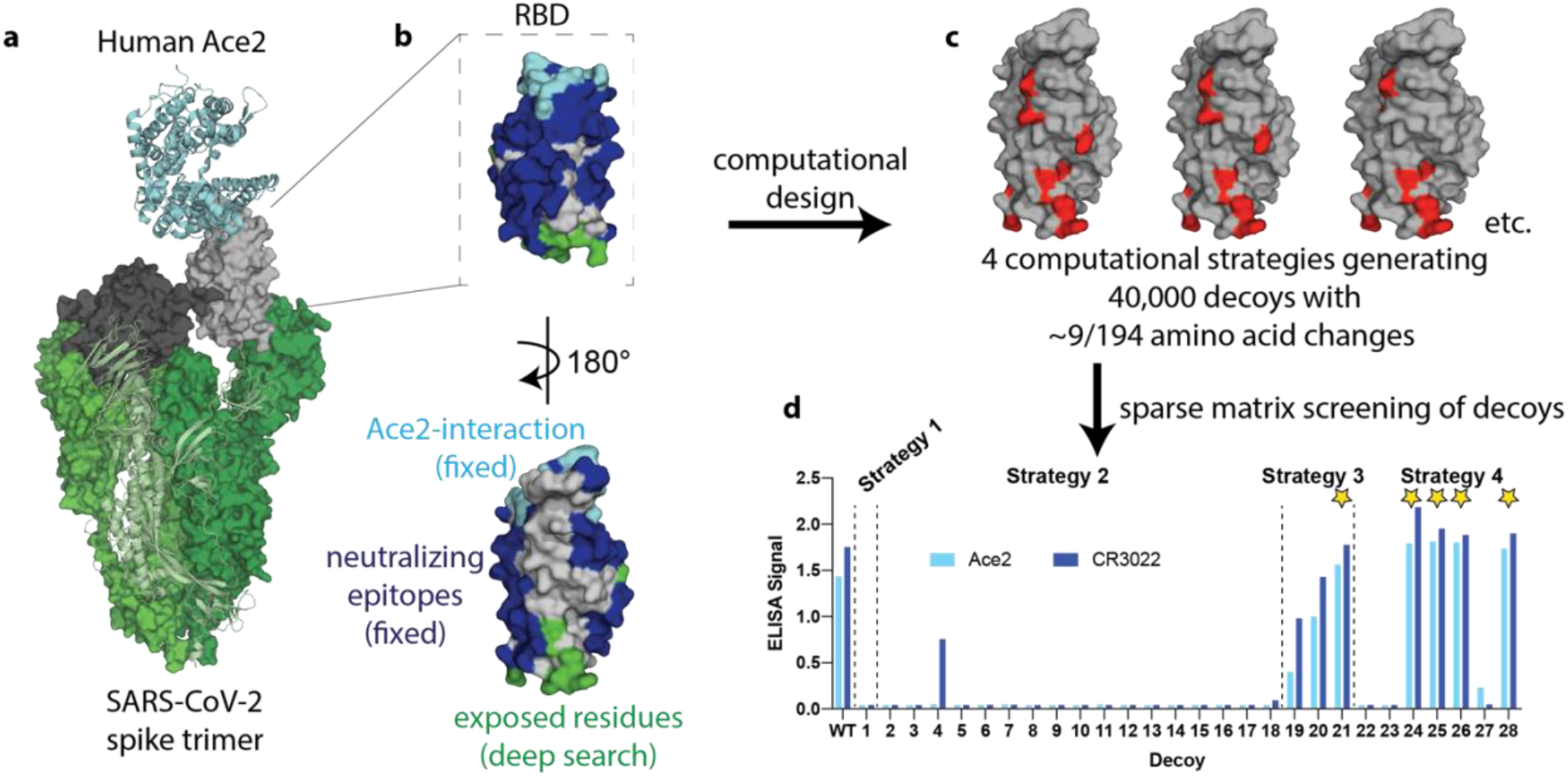
Overview of SPEEDesign pipeline used to create RBD immunogens. **a**, The SARS-CoV-2 spike trimer (green) binds human Ace2 (cyan) to mediate viral entry. This interaction is mediated by the “up” conformation of the RBD (grey), which can also exist in a down conformation (black). **b**, The RBD design process retained the Ace2-interaction surface (cyan) and all known SARS neutralizing epitopes (blue). Residues exposed upon isolation of the RBD (green) were heavily designed while all other residues (grey) were designed more conservatively. **c**, Four computational strategies were used to create 40,000 decoys, each of which has an average of nine amino acid changes (red). **d**, 28 sequences sampling the top scoring decoys were screened *in vitro*, identifying 5 lead immunogen candidates (stars).

## Results

### SPEEDesign of novel RBD immunogens

We developed a novel computational design and *in vitro* screening pipeline termed Stabilizer for Protein Expression and Epitope Design (SPEEDesign) to design improved RBD vaccine candidates. The objectives of SPEEDesign are to improve the protective efficacy of a given antigen by: 1) focusing the immune response towards potently neutralizing antibody epitopes on the antigen; 2) reducing or eliminating the immune responses to poorly neutralizing and/or immunodominant epitopes within the antigen; 3) optimizing the thermal stability of the antigen to increase its durability *in vivo* following immunization; 4) promoting conformational states of a protein antigen that may be hidden, for example the “up” state of the RBD in the spike protein. These four objectives are achieved by careful definition of the role of individual amino acids within the antigen, and by extension their mutability in ROSETTA design strategies coupled to a clustering and *in vitro* screening approach.

The Ace2 interface and all known neutralizing epitopes within the RBD were retained and not allowed to vary, as the designed immunogen is expected to focus the immune response to these segments (Fig. 1b). Residues that are exposed in the RBD “up” conformation, or that are exposed upon extraction of the RBD from the full-length (FL)-spike trimer, were thoroughly searched during the design process to identify amino acid changes that would stabilize an accessible RBD. All remaining residues, which likely include surface residues that comprise poorly or non-neutralizing epitopes, were allowed to sample a limited sequence space defined by energetic and evolutionary restraints^21^.

Four different computational ROSETTA design strategies, which differ in the depth of design for each class of residues, were used to identify amino acid changes that would achieve the goals of SPEEDesign (Fig. 1c). Ten thousand decoys were produced for each computational strategy, each of which had approximately nine amino acid changes from the native RBD sequence. Twenty-eight decoys were selected using a clustering algorithm designed to broadly sample the 40,000 computational decoys (Fig. 1d). These decoys are distributed between strategies according to the sequence diversity produced by each strategy. For example, more decoys were selected from strategy 2 than strategy 1 because strategy 2 samples a much larger sequence space.

The 28 representative sequences were screened *in vitro* for expression and presentation of neutralizing epitopes (Fig. 1d). We used Ace2 to probe the integrity of the most potent neutralizing epitope and the monoclonal antibody CR3022, which recognizes an orthogonal surface conserved between coronaviruses. Interestingly, all successful lead immunogen candidates derive from strategies 3 and 4, which sample a moderate sequence space, rather than the highly restricted strategy 1 or highly divergent strategy 2. Additionally, the immunogen with the best ROSETTA score from each strategy (decoys 1, 2, 19 and 22) were not successful, indicating that ROSETTA score alone is not a suitable predictor for a successful design. The clustering and screening strategies unique to the SPEEDesign pipeline are a critical advancement that allows the identification of improved immunogens among the many decoys produced during ROSETTA design.

### Designed immunogens have increased expression and stability

Five lead candidate immunogens (**1**–**5**, corresponding to decoys 28, 24, 25, 26, and 21, respectively) were expressed and purified to determine their biophysical characteristics. All immunogens are well-folded monomeric proteins that can be highly purified with yields much greater than native wild type (WT) RBD (Fig. 2a, b, and d). Three immunogens (**1**, **2**, and **3**) can be purified with final yields ≥200 mg/L, or more than 4-fold higher than WT RBD (Fig. 2d). Furthermore, these yields are approximately 6-fold higher than the yields reported for an optimized FL-spike ectodomain known as “hexapro” and 200-fold higher than the well-established 2P prefusion-stabilized FL-spike ectodomain, or expressed as a per mole or per epitope equivalent these improvements are 35 and 1,200 fold higher^22^. These yields were achieved without optimization of the expression and purification system, suggesting that process development could reach production levels sufficient for the development of a highly cost-effective vaccine.

**Fig. 2.**
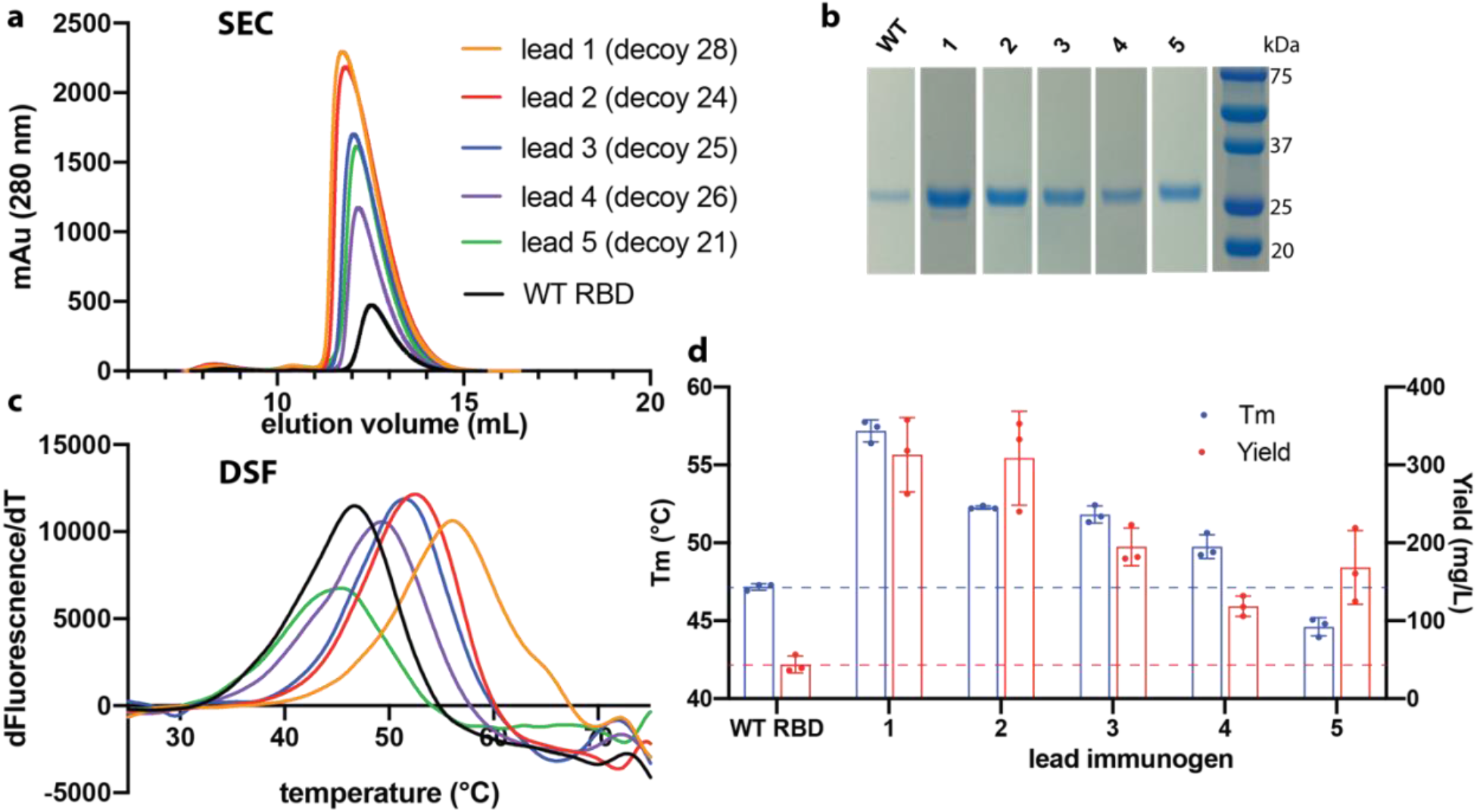
Stability and yield are higher for immunogens than WT RBD. **a**, All five immunogens express at higher levels than WT RBD and elute as monomers by size-exclusion chromatography. **b**, SDS-PAGE confirms the high purity and yield of RBD immunogens. **c**, Differential scanning fluorimetry indicates that 4 immunogens have higher thermostability than WT RBD. **d**, T_m_ and purification yield averages and standard deviations from three separate purifications.

The thermostability of four lead immunogens was also higher than WT RBD (Fig. 2c and d). One immunogen (**1**) has a melting temperature (T_m_) of 57 °C, or 10 °C higher than WT RBD, and two additional immunogens (**2** and **3**) have greater than 5 °C increases in T_m_. Expression yield roughly correlates with T_m_, and immunogens **1**, **2**, and **3** have substantially higher yields and stability than WT RBD. Increased thermostability likely contributes to the increase in immunogen yield, will likely increase the half-life of the antigen in the body, and will likely improve stability during storage, transportation, and administration of a vaccine, alleviating cold-chain requirements.

### The structural basis for enhanced immunogen stability

The five lead immunogens have a distinct pattern of amino acid changes that provide a mechanistic basis for their enhanced biophysical characteristics. There are nine positions at which multiple lead immunogens have non-conservative amino acid substitutions relative to the native protein sequence (Fig. 3a and Extended Data Table 1). At four of these positions, the immunogens share similar amino acid identities: 1) Alanine 363 is changed to a large hydrophobic residue in all five lead immunogens, 2) I468T in all five lead immunogens, 3) H519D in all five lead immunogens, 4) A522P in 3/5 lead immunogens.

**Fig. 3.**
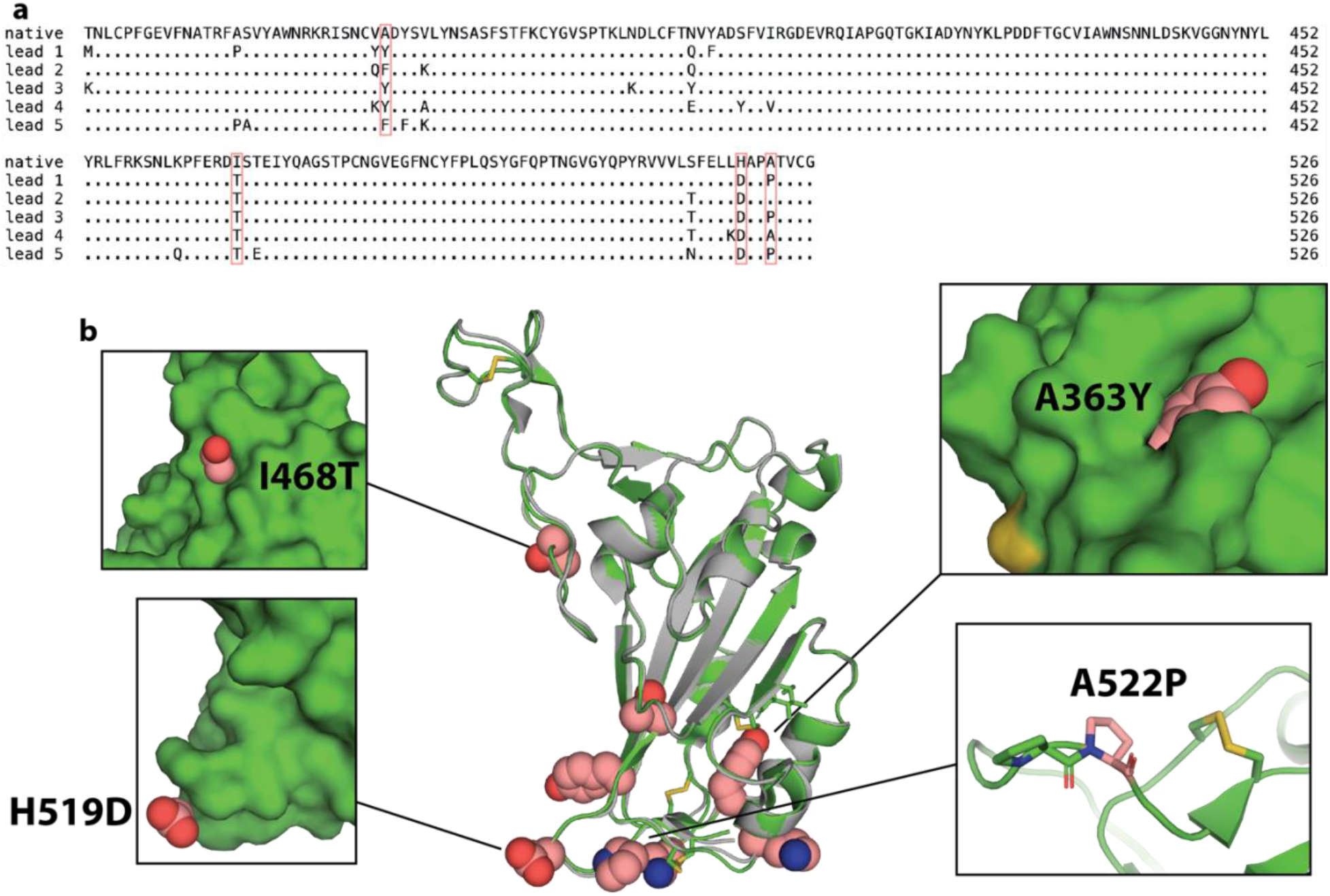
Structural basis for immunogen stabilization. **a**, Sequence alignment of amino acid changes in lead immunogens relative to the native RBD sequence. Four recurring changes are highlighted in pink. **b**, The crystal structure of immunogen **3** (green) is globally similar to WT RBD (grey; PDB:7BWJ) despite eight amino acid changes (pink spheres). Insets illustrate key substitutions.

To determine how these amino acid changes improve the stability of the immunogens, we solved the crystal structure of lead immunogen **3** in complex with the neutralizing Fab P2B-2F6 (Extended Data Fig. 1a)^23^. The overall structure of immunogen **3** is very similar to the native RBD with a Cα RMSD of only 0.476 Å across all residues, indicating that the amino acid changes do not alter the overall shape, secondary structure, or tertiary structure of the RBD (Fig. 3b and Extended Data Table 2). The enhanced biophysical characteristics are driven by local structural changes around substituted side-chains. For example, A363Y is a space-filling substitution that likely stabilizes the protein fold and proximal di-sulfide bonds. Ile468 and His519 are buried in the RBD-down conformation and become exposed in the RBD-up conformation (Extended Data Fig. 2). Therefore, the I468T and H519D substitutions increase the hydrophilicity of solvent-exposed residues, likely improving the solubility of RBD immunogens and potentially promoting the RBD-up conformation in the context of the spike trimer. Finally, A522P creates a tandem proline sequence that likely promotes a sharp kink in the backbone adjacent to a disulfide bond. All amino acid changes in the immunogens, including the 4 key positions, are distal to the changes found in naturally occurring variants of concern (*e.g.*, B.1.17, P.1, and B.1.351), suggesting that these immunogens are compatible with next-generation vaccines that derive from variant sequences (Extended Data Fig. 1b and c). Indeed, immunogens **1** – **3** increase expression in the context of the B.1.351 variant to an even greater extent than the original variant (Extended Data Fig. 1d). Thus, the stabilizing mutations promote the stability of the immunogens through diverse structural and energetic mechanisms that do not impact the global antigen structure and that are compatible with next-generation vaccines.

### Neutralizing epitopes are unperturbed on designed immunogens

We probed the Ace2 receptor-binding site and several key three-dimensional antibody epitopes to establish that the stabilizing mutations did not disrupt key interacting residues or neutralizing epitopes in the immunogens (Extended Data Fig. 3). Ace2 binds to the end of the RBD, as do potently neutralizing Ace2-blocking antibodies that include the mAbs REGN10933 and P2B-2F6^9,23,24^. Additional neutralizing mAbs CR3022 and S309 bind to opposing sides to Ace2 and do not overlap with each other or the Ace2 binding site (Extended Data Fig. 3a)^25,26^. The Ace2 binding site and these four epitopes therefore report the structural integrity of almost all neutralizing surfaces retained during the design process. The designed immunogens bind Ace2, REGN10933, P2B-2F6, CR3022 and S309 at least as well as WT, with the exception of immunogen **5** showing a slight decrease in binding to P2B-2F6, suggesting that the stabilizing changes made to the immunogens **1-4** do not perturb nearby surfaces and critical neutralizing epitopes (Extended Data Fig. 3b). We further used biolayer interferometry (BLI) to measure the integrity of the Ace2 binding site and REGN10933 epitope in a more quantitative fashion (Extended Data Fig. 3c). Consistent with the ELISA results, immunogens **1** - **4** bind both probes with biophysical parameters very similar to WT RBD (Extended Data Table 3). The Ace2, REGN10933, P2B-2F6, CR3022 and S309 probes used here represent all four classes of epitopes recognized by known RBD antibodies^27^. Therefore, neutralizing epitopes are unperturbed on RBD immunogens **1** - **4** and an immune response to an immunogen would be expected to recognize the native RBD protein and SARS-CoV-2 virus.

### Vaccination with immunogens enhances antibody neutralization

Mice were immunized with the five lead immunogens to determine if the amino acid changes improve the neutralizing antibody response (Fig. 4). CD-1 outbred mice were used to mimic the genetic diversity found in the human population more closely than inbred mice. All immunogens generate antibodies that recognize trimeric FL-spike ectodomain and **1**, **2**, and **3** generate geometric mean titers (GMT) greater than WT RBD, consistent with their enhanced biophysical characteristics (Fig. 4b). We measured inhibition of RBD/Ace2 binding to evaluate the titers of functional antibodies elicited by each antigen (Fig. 4c). Again, **1**, **2**, and **3** outperform WT RBD, generating Ace2-blocking GMTs up to 10-fold higher. We also measured neutralizing titers in a pseudoviral neutralization assay and again found that **1, 2**, and **3** outperform WT RBD (Fig. 4d). Immunogen **3** elicits neutralizing titers significantly greater than WT RBD with a GMT more than 30-fold higher than WT, an even greater enhancement than the ~10-fold increase provided by the 2P stabilizing mutation^6^.

**Fig. 4.**
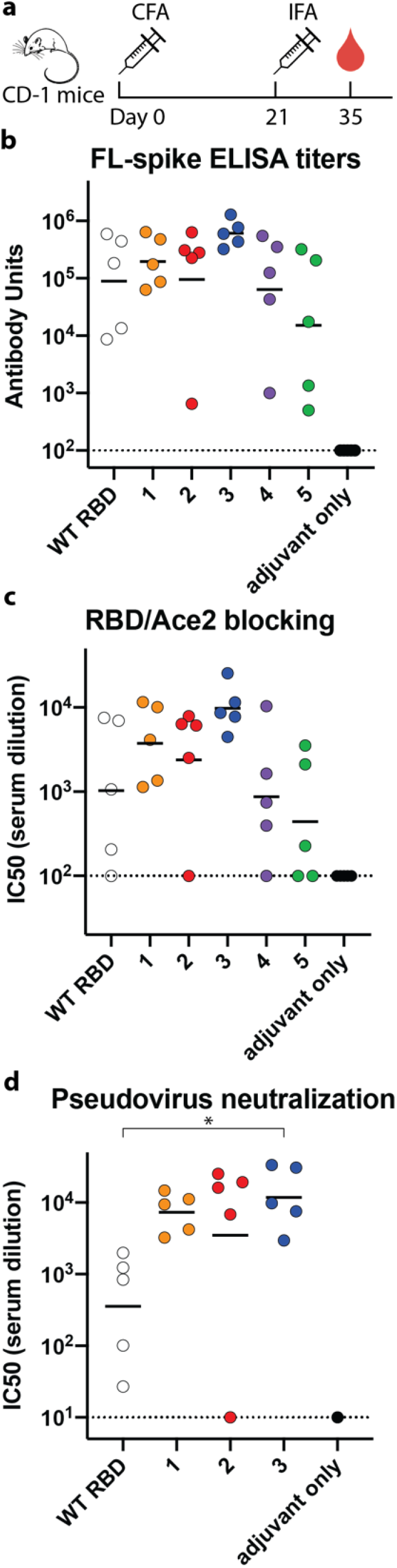
Blocking antibody titers are higher in mice immunized with immunogens than WT RBD. **a**, Immunization and blood draw schedule for CD-1 mice. **b**, Serum ELISA titers against trimeric FL-spike ectodomain. Dashed line indicates detection limit of assay and bars represent geometric mean titers (GMT). **c**, Titers of antibodies blocking Ace2/RBD interaction depicted as described in **b**. **d**, Pseudovirus neutralization titers depicted as described in **b**. Statistical comparisons were made using a Kruskal-Wallis ANOVA followed by Dunn’s comparison with WT RBD (* = p < 0.05). p-values are 0.21, 0.14, 0.043 for **1**, **2**, and **3**, respectively.

## Discussion

These results demonstrate that a handful of amino acid changes to the RBD can dramatically improve the protective antibody response. A 30-fold increase in neutralizing antibody titers is equivalent to lengthening the lifetime of neutralizing antibodies by approximately five half-lives of decay, suggesting that immunogens would confer a much longer duration of antibody-mediated protection than WT RBD. Additionally, this 30-fold enhancement is greater than the 1.5-12 fold reduction in neutralizing potential against new SARS-CoV-2 variants, such as B.1.351, suggesting that these immunogens could increase the breadth of protection, in addition to the duration^28–31^.

The three lead immunogens appear to have achieved the objectives set out for SPEEDesign. While it is difficult to pinpoint the cause of improved vaccine performance, it is interesting to note that immunogen **3** elicited 30-fold greater neutralizing antibody titers, but only 7-fold greater Spike ELISA titers. This indicates that SPEEDesign achieved the major objectives of focusing the immune response to potently neutralizing epitopes and away from poorly neutralizing epitopes as the increase in spike titers alone is insufficient to explain the large improvement in neutralizing antibody titers. We expect that SPEEDesign will achieve similar success with other antigens that have structurally characterized neutralizing epitopes and will be useful for designing vaccines against many other pathogens.

These monomeric designed RBD immunogens are expected to be effective vaccines in their own right. In addition to the enhanced protection, the improvements in stability and yield suggest that recombinant RBD immunogens would be easily manufactured and distributed to meet the global need for a SARS-CoV-2 vaccine. These manufacturing and distribution benefits would be even more meaningful in the event that seasonal updates and mass vaccination campaigns are required. Finally, these immunogens can be easily improved by established methods of foldon trimerization, multimerization, and/or nanoparticle display^14,32–35^.

The designed immunogens can also be easily adapted to all other vaccine platforms by making amino acid changes to the RBD sequence within the FL-spike or within RBD-only vaccines. The changes identified here do not exclude the existing improvements in spike, they are fully compatible with changes in the S2 domain, and they are compatible with all known changes in emerging SARS-CoV-2 variants of concern. The SPEEDesign changes stabilize the RBD in isolation and may provide additional benefit to a FL-spike vaccine by restricting the down conformation of the RBD. Regardless of the platform, these amino acid changes would be expected to increase RBD stability, enhance the protective immune response, and lengthen the lifetime and breadth of protection conferred by a SARS-CoV-2 vaccine.

## Online Methods

### SPEEDesign in vitro screening

Synthetic DNA coding for secreted RBD immunogens was cloned (GenScript) into a customized pHL-sec expression plasmid. pHL-sec was a gift from Edith Yvonne Jones (Addgene plasmid # 99845; http://n2t.net/addgene:99845; RRID:Addgene_99845)^45^. Plasmid was transfected into human expi293F cells and grown in a 96-well plate according to manufacturer instructions (ThermoFisher Scientific). Cell-free supernatant was harvested after 5 days of expression.

Cell-free supernatant was diluted in PBST + 2% BSA and added to Ni-NTA HisSorb Plates (Qiagen) to capture His-tagged immunogens. After incubation for 1 hour at room temperature, plates were washed three times with PBST. Neutralizing epitopes were probed using an Ace2-Fc(IgG1) fusion (0.2 ug/well) or a human IgG1 antibody containing the CR3022 variable domain (0.05 ug/well). After incubation for 1 hour at room temperature, plates were washed three times with PBST and 200 μl 1:5000 peroxidase-conjugated anti-human IgG was added (Jackson ImmunoResearch Laboratories, Inc. Cat.# 109-035-098). Plates were incubated 30 minutes at room temperature and washed three times with PBST. Finally, 70 μl Tetramethylbenzidine (TMB) (MilliporeSigma) was added and incubated 5 minutes at room temperature before quenching with 70 μl 2 M H_2_SO_4_. Absorbance at 450 nm was measured using a Biotek Synergy H1 plate reader.

### Immunogen expression, purification, and calculation of yields

Recombinant RBD immunogens were expressed in expi293F cells, as described for SPEEDesign *in vitro* screening above. Trimeric FL-spike ectodomain was also expressed in expi293F cells using a modified pHL-sec vector. The construct contains amino acids 16-1208 of the spike protein followed by a foldon trimerization domain and a 6-His tag. This construct also contains the “2P” stabilizing mutations at K986P and V987P and mutation of the furin cleavage site (682-685 GSAS to RRAR)^19^.

Cell-free supernatant was harvested after 4 days and His-tagged immunogens were purified by gravity chromatography using Ni Sepharose excel resin according to manufacturer instructions (Cytiva). Immunogens were further purified by size-exclusion chromatography using a Superdex 75 Increase 10/300 GL column (RBD immunogens) or Superose 6 Increase 10/300 GL column (trimeric FL-spike) equilibrated in 1x PBS. Fractions corresponding to trimeric FL-spike or monomeric RBD were pooled, snap frozen in liquid nitrogen, and stored at −80 °C.

Transfection, expression, and purification was performed in triplicate on three separate days to calculate RBD immunogen purification yields. Each replicate consisted of a 30 mL culture, and yields were calculated by integrating the area under the monomeric peak on the Abs_280_ chromatogram during size-exclusion chromatography. These yields closely matched yields calculated from pooled fractions. Extinction coefficients were calculated using the ExPASy ProtParam tool^46^.

### Antibody expression and purification

Antibodies for ELISAs were created by fusing the variable regions for the indicated antibody to the human IGHG*01, IGKC*01, or IGLC2*02 constant regions and cloning into the pHL-sec plasmid (GenScript). The Ace2-Fc fusion was expressed from pcDNA3: pcDNA3-sACE2(WT)-Fc(IgG1) was a gift from Erik Procko (Addgene plasmid # 145163; http://n2t.net/addgene:145163; RRID:Addgene_145163)^47^. Heavy and light chain plasmids were mixed in equal amounts and transfected into expi293F cells according to manufacturer instructions and cell-free supernatant was harvested after 4 days of expression (ThermoFisher Scientific).

Cell-free supernatant was batch incubated with protein A agarose resin (GoldBio) for 1 hour at room temperature. Resin was collected and washed with 10 column volumes (CV) protein A IgG binding buffer (ThermoFisher Scientific). Protein was eluted with 10 CV IgG elution buffer (ThermoFisher Scientific) and neutralized with 1 CV 1 M Tris pH 9.0. Antibodies were concentrated and buffer exchanged into PBS using an amicon centrifugal filter (MilliporeSigma). Ace2-Fc was further purified by size-exclusion chromatography using a superdex 200 increase 10/300 GL column (Cytiva) equilibrated in 1x PBS.

### Differential Scanning Fluorimetry

DSF was performed using the Protein Thermal Shift Dye Kit according to manufacturer instructions (ThermoFisher Scientific). Final reactions contained 0.125 mg/mL purified immunogen, 1x Protein Thermal Shift buffer, 1x Thermal Shift Dye, and 0.63x PBS. Fluorescence was monitored using a 7500 Fast Real-Time PCR system (ThermoFisher Scientific) as the temperature was increased from 25 °C to 95 °C at a ramp rate of 1%. Melting temperature (T_m_) was calculated as the peak of the derivative of the melt curve. DSF reactions were performed in technical quadruplicate on each plate and in biological triplicate using three different protein preps on 3 separate days. Technical replicates were averaged to calculate the T_m_ for a biological replicate, and the three biological replicates were averaged to calculate the reported T_m_.

### Crystallization

The antigen-binding fragment (Fab) of P2B-2F6 was used to promote crystallization of lead immunogen **3**. P2B-2F6 Fab was produced by fusing the P2B-2F6 variable region to a His-tagged human IGHG*01 C_H_1 domain and cloning into pHL-sec (GenScript). Heavy chain Fab plasmid was co-transfected 1:1 with light chain in expi293F cells according to manufacturer instructions and cell-free supernatant was harvested after 4 days of expression (ThermoFisher Scientific). His-tagged Fab was purified from cell-free supernatant by gravity chromatography using Ni Sepharose excel resin according to manufacturer instructions (Cytiva). Fab was further purified by size-exclusion chromatography using a Superdex 75 increase 10/300 GL column (Cytiva) equilibrated in 1x PBS.

Purified Fab was mixed with purified lead immunogen **3** in a 1.5:1 ratio and incubated 30 minutes on ice. Complex was purified by size-exclusion chromatography on a Superdex 200 increase 10/300 GL column equilibrated in 10 mM Na-HEPES pH 7.4, 100 mM NaCl. Purified complex was concentrated to 18 mg/mL using an amicon centrifugal filter (MilliporeSigma) and crystal trays were setup using a mosquito crystal robot (STP Labtech). Drops contained 0.2 μl complex and 0.2 μl reservoir solution (0.2 M sodium fluoride, 20% w/v PEG 3,350). Crystals were grown by hanging-drop vapor diffusion at 18 °C for 13 days. Crystals were cryoprotected in a solution that contained 7 uL well-solution and 3 μl 100% glycerol and flash frozen in liquid nitrogen.

### Data collection and structure determination

Crystal diffraction data were collected at 1.033 Å at 100 °K on the GM/CA 23-ID-D beamline at the Advanced Photon Source. Data were processed using autoPROC^48^. Reflections were indexed and integrated using XDS^49^. Data were scaled and merged using AIMLESS^50,51^. The P2B-2F6/WT RBD structure (PDB: 7BWJ) was used as a starting model for rigid body refinement in PHENIX Refine^52^. This model was then edited in COOT to incorporate the amino acid changes present in immunogen **3** and subsequent rounds of refinement and model building were performed with COOT and PHENIX Refine^53^. The final model was evaluated with MolProbity, which showed good geometry, with 96.0% of the residues as Ramachandran favored and 0% outlier residues (Extended Data Table 2)^54^. Software used in this project was curated by SBGrid^55^.

### ELISA analysis of immunogen epitopes

Nunc MaxiSorp plates (ThermoFisher Scientific) were coated with 100 μl 0.01 mg/mL purified immunogen diluted in 50 mM Na-carbonate pH 9.5. Plates were coated overnight at 4 °C then washed three times with PBST. Plates were blocked 1 hour at room temperature with 2% BSA in PBST then washed three times with PBST. 100 μl primary antibody was added to each well at the indicated concentration: Ace2 – 3.1 ng/mL, REGN10933, CR3022, and S309 – 1.5 ng/mL, and P2B-2F6 – 7.5 ng/mL. Primary antibody was incubated 1 hour at room temperature then plates were washed three times with PBST and 200 μl 1:5000 peroxidase-conjugated anti-human IgG was added (Jackson ImmunoResearch Laboratories, Inc. Cat.# 109-035-098). Plates were incubated 30 minutes at room temperature and washed three times with PBST. Finally, 70 μl Tetramethylbenzidine (TMB) (MilliporeSigma) was added and incubated 10 minutes at room temperature before quenching with 70 μl 2 M H_2_SO_4_. Absorbance at 450 nm was measured using a Biotek Synergy H1 plate reader.

### Biolayer interferometry

The binding affinity of purified immunogens to REGN10933 and Ace2-Fc was measured using a kinetic BLI assay using an Octet-Red96e (Sartorius). REGN10933 IgG or Ace2-Fc was buffer exchanged into HBS-EP buffer (10 mM Na-HEPES pH 7.4, 150 mM NaCl, 3 mM EDTA, 0.005% v/v P20 surfactant) using Zeba spin desalting columns (ThermoFisher Scientific). REGN10933 or Ace2-Fc was loaded onto Anti-hIgG Fc Capture (AHC) biosensors (Sartorius) over the course of 300 seconds, until reaching a signal of ~0.6 nm. BLI pins were then immersed in immunogens 2-fold serially diluted in HBS-EP buffer (150 nM to 2.34 nM). After 300 seconds, pins were immersed in HBS-EP buffer to measure dissociation. Association rate (k_a_), dissociation rate (k_dis_), and dissociation constant (K_D_) were globally fit using a 1:1 binding model in Data Analysis HT 12.0 (Sartorius). Three independent protein preps (biological replicates) were each measured in technical triplicate. Values reported are the average and standard deviation between biological replicates.

### Mouse immunizations

Mouse immunogenicity studies were performed under the guidelines and approval of the Institutional Animal Care and Use Committee (IACUC) at the National Institutes of Health. Five 5-6 week old female CD-1 mice (Charles River Laboratories) per group were immunized with 10 μg antigen each. Antigen was formulated as a 1:1 ratio in Complete Freund’s Adjuvant (MilliporeSigma) on day 0 and Incomplete Freund’s Adjuvant (MilliporeSigma) on day 21 and 100 μl formulated antigen was delivered by intraperitoneal injection. Blood was collected on day 35 and serum was separated and stored at −80 °C.

### Serum antibody titer ELISA

Nunc MaxiSorp plates (ThermoFisher Scientific) were coated with 100 μl 0.01 mg/mL purified trimeric FL-spike ectodomain diluted in 50 mM Na-carbonate pH 9.5. Plates were incubated overnight at 4 °C then washed three times with PBST. Plates were blocked 1 hour at room temperature with 2% BSA in PBST then washed three times with PBST. Serum was diluted in 2% BSA in PBST and 100 μl was added to each well. After 1 hour incubation at room temperature, plates were washed three times with PBST and 200 μl 1:5000 peroxidase-conjugated anti-mouse IgG was added (Jackson ImmunoResearch Laboratories, Inc. Cat.# 115-035-164). Plates were incubated 30 minutes at room temperature and washed three times with PBST. Finally, 70 μl Tetramethylbenzidine (TMB) (MilliporeSigma) was added and incubated 20 minutes at room temperature before quenching with 70 μl 2 M H_2_SO_4_. Absorbance at 450 nm was measured using a Biotek Synergy H1 plate reader.

Pooled serum from mice immunized with WT RBD was used as a standard curve on each plate to calculate the antibody titers of individual animals in all groups. One antibody unit (AU) was defined as the dilution of the standard serum required to achieve an Abs_450_ value of 1. Each plate included triplicate 2-fold serial dilutions of the standard serum from 20 to 0.01 AU. Serum from each animal was diluted such that the Abs_450_ fell in the informative portion of the standard curve between 0.1 and 2.0. The Abs_450_ values for the standard curve were fit to a 4-parameter logistic curve, which was used to convert Abs_450_ values to AU for each individual animal. AU values for each individual animal were measured in triplicate on separate plates and the average is reported.

### RBD/Ace2 blocking assay

Nunc MaxiSorp plates (ThermoFisher Scientific) were coated with 100 μl 0.32 μg/mL purified WT RBD diluted in 50 mM Na-carbonate pH 9.5. Plates were incubated overnight at 4 °C then washed three times with PBST. Plates were blocked 1 hour at room temperature with 2% BSA in PBST then washed three times with PBST. Serum was diluted in 2% BSA in PBST in a 3-fold dilution series from 1:100 to 1:218,700. 50 μl serum was mixed with 10 μl 30 nM Ace2-Fc or 10 μl buffer as a background control. 50 μl serum mixture was added to RBD-coated plates and incubated 1 hour at room temperature. Plates were washed three times with PBST and 200 μl 1:30,000 peroxidase-conjugated anti-human IgG was added (Jackson ImmunoResearch Laboratories, Inc. Cat.# 109-035-098). Plates were incubated 30 minutes at room temperature and washed three times with PBST. Finally, 70 μl Tetramethylbenzidine (TMB) (MilliporeSigma) was added and incubated 20 minutes at room temperature before quenching with 70 μl 2 M H_2_SO_4_. Absorbance at 450 nm was measured using a Biotek Synergy H1 plate reader.

Ace2/RBD binding inhibition was calculated by first subtracting the Abs_450_ values of the background controls lacking Ace2-Fc. Eight wells without serum were used to calculate the maximum signal. Inhibition was calculated using the following formula:

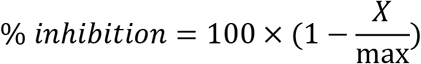

where X is the Abs_450_ of a well after background-subtraction and max is the average of the 8 samples without serum after background-subtraction.

% inhibition values were measured for each serum dilution in triplicate and average values were plotted in GraphPad Prism 8. Data were fit using a normalized dose response curve with a variable slope:

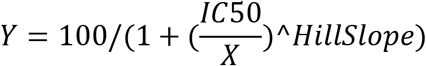

where X is the serum dilution, Y is the % inhibition, and HillSlope and IC50 are calculated parameters corresponding to the slope of the curve and the dilution at which 50% inhibition occurs, respectively.

IC50 values for each animal were log-transformed and plotted, along with the geometric mean value for each group.

### Pseudoviral neutralization

Mouse serum was diluted in a duplicate 4-fold series from 1:5 to 1:81,920. Serum was mixed 1:1 with pseudovirus containing the SARS-CoV-2 spike protein and a luciferase reporter in a 96-well plate (GenScript). After 1 hour incubation at room temperature, 20,000 HEK293 cells overexpressing Ace2 were added to each well. Cells and pseudovirus were incubated at 37 °C, 5% CO2 for 48 hours, after which culture medium was removed and Bio-Glo luciferase reagent (Promega) was added to the wells. Luciferase signal was measured using an EnVision plate reader and %inhibition was calculated using the following formula:

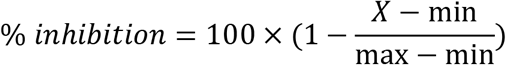

Where X is the luciferase signal, min is the average signal from duplicate wells without pseudovirus, and max is the average signal from duplicate wells without serum.

log IC50 values were calculated in the same manner described for the RBD/Ace2 blocking assay.

## Acknowledgments

We thank all the members of LMIV COVID group for discussions of results and Nathan Max and Palak Patel for experimental assistance. This work was funded by the Intramural Research Program of the National Institute of Allergy and Infectious Diseases (NIAID), National Institutes of Health. This study used the Office of Cyber Infrastructure and Computational Biology (OCICB) High Performance Computing (HPC) cluster at the National Institute of Allergy and Infectious Diseases (NIAID), Bethesda, MD. GM/CA@APS has been funded in whole or in part with Federal funds from the National Cancer Institute (ACB-12002) and the National Institute of General Medical Sciences (AGM-12006). This research used resources of the Advanced Photon Source, a U.S. Department of Energy (DOE) Office of Science User Facility operated for the DOE Office of Science by Argonne National Laboratory under Contract No. DE-AC02-06CH11357.

## Author contributions

N.H.T. conceptualized the design procedure and application to COVID-19, analyzed data, acquired funding, wrote the manuscript, developed software, and supervised the project; THD conceptualized the application to COVID-19, curated data, analyzed data, contributed to funding applications, wrote the manuscript, developed experimental methods, and performed experiments; W.K.T. performed experiments, created figures, and edited the manuscript; B.B. performed experiments; S.O. performed experiments; T.O. performed experiments; N.D.S. managed animals studies and edited the manuscript; L.E.L. managed animal studies.

## Materials & Correspondence

N.H.T.

## Extended Data

**Extended Data Fig. 1.**
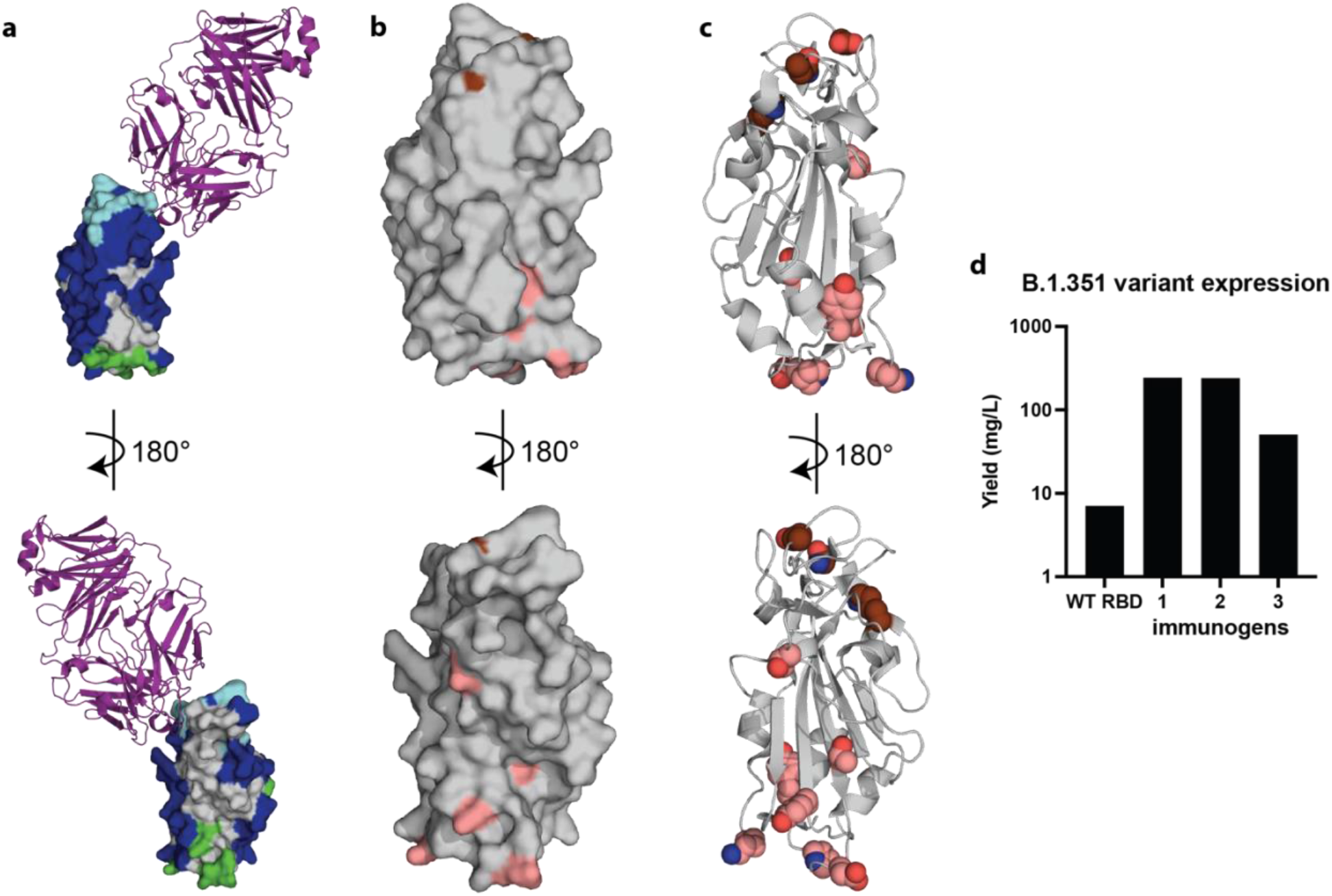
The amino acid changes in immunogen 3 are compatible with variants of concern. **a**, Crystal structure of P2B-2F6 Fab (purple) bound to immunogen 3 coloured as in Fig. 1 (cyan: Ace2 contact; blue: neutralizing epitopes; grey: intermediate sequence design depth; green: exposed residues that were heavily designed) **b, c** Immunogen 3 with beneficial amino acid changes (pink) and mutations in variants of concern (brown) (K417, E484, and N501) as surface (**b)** and cartoon (**c**) representation. **d**, Purification yields for proteins containing the B.1.351 variant amino acid changes.

**Extended Data Fig. 2.**
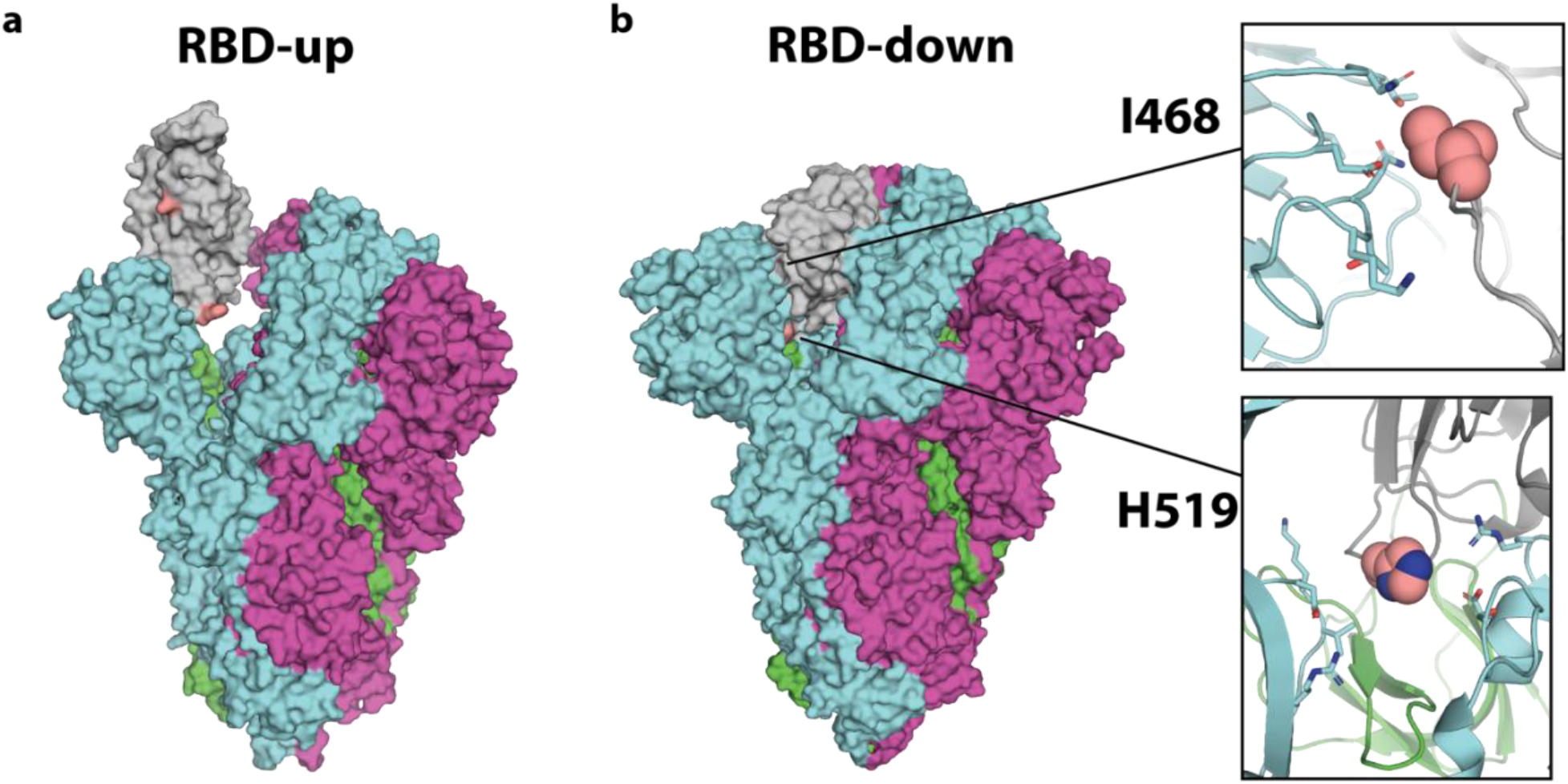
I468 and H519 are buried in the RBD-down conformation. **a**, I468 and H519 (pink) are solvent exposed when the RBD (grey) is in the up conformation (PDB 6VSB and 6MOJ)^24,56^. Protomers are shown in green, cyan, and magenta with the RBD of the green protomer in the up conformation. **b**, I468 and H519 (pink) become buried in the closed conformation where they make contacts with the neighboring protomer (cyan) (PDB 6XLU). Mutation of these residues to more hydrophilic residues could promote the RBD-up conformation.

**Extended Data Fig. 3.**
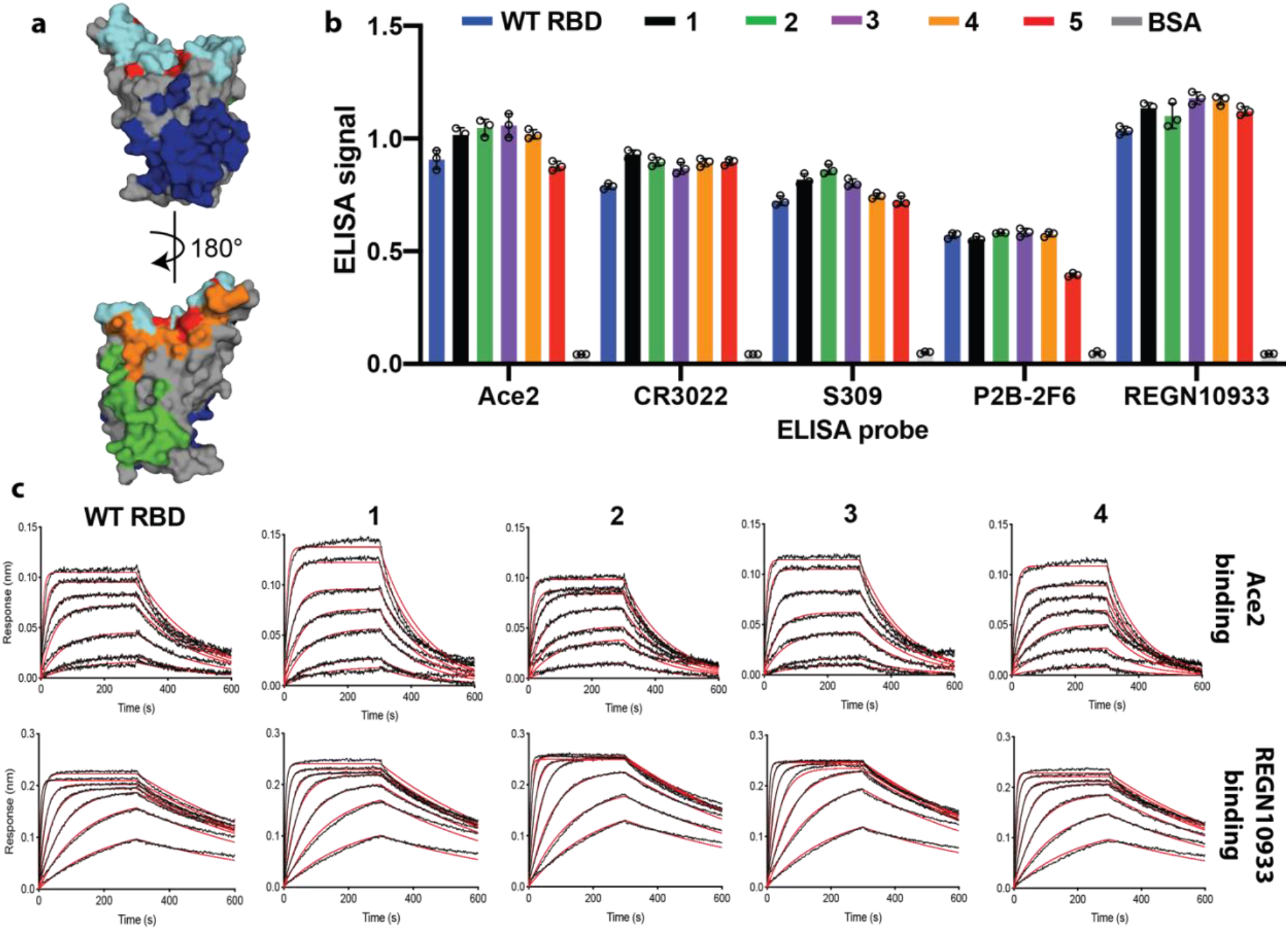
Neutralizing epitopes are unperturbed on RBD immunogens. **a**, Five distinct three-dimensional neutralizing epitopes covering the majority of the protein surface were probed for each immunogen (Ace2:cyan, REGN10933:red, P2B-2F6:orange, S309:green, CR3022:blue). **b**, ELISA probes bind to all five epitopes on the immunogens **c**, Representative BLI traces used to quantitatively measure the binding of the immunogens to two probes, demonstrating the high integrity of these epitopes. Immunogen concentrations begin at 150 nM and decrease in 2-fold increments.

**Extended Data Table 1.**
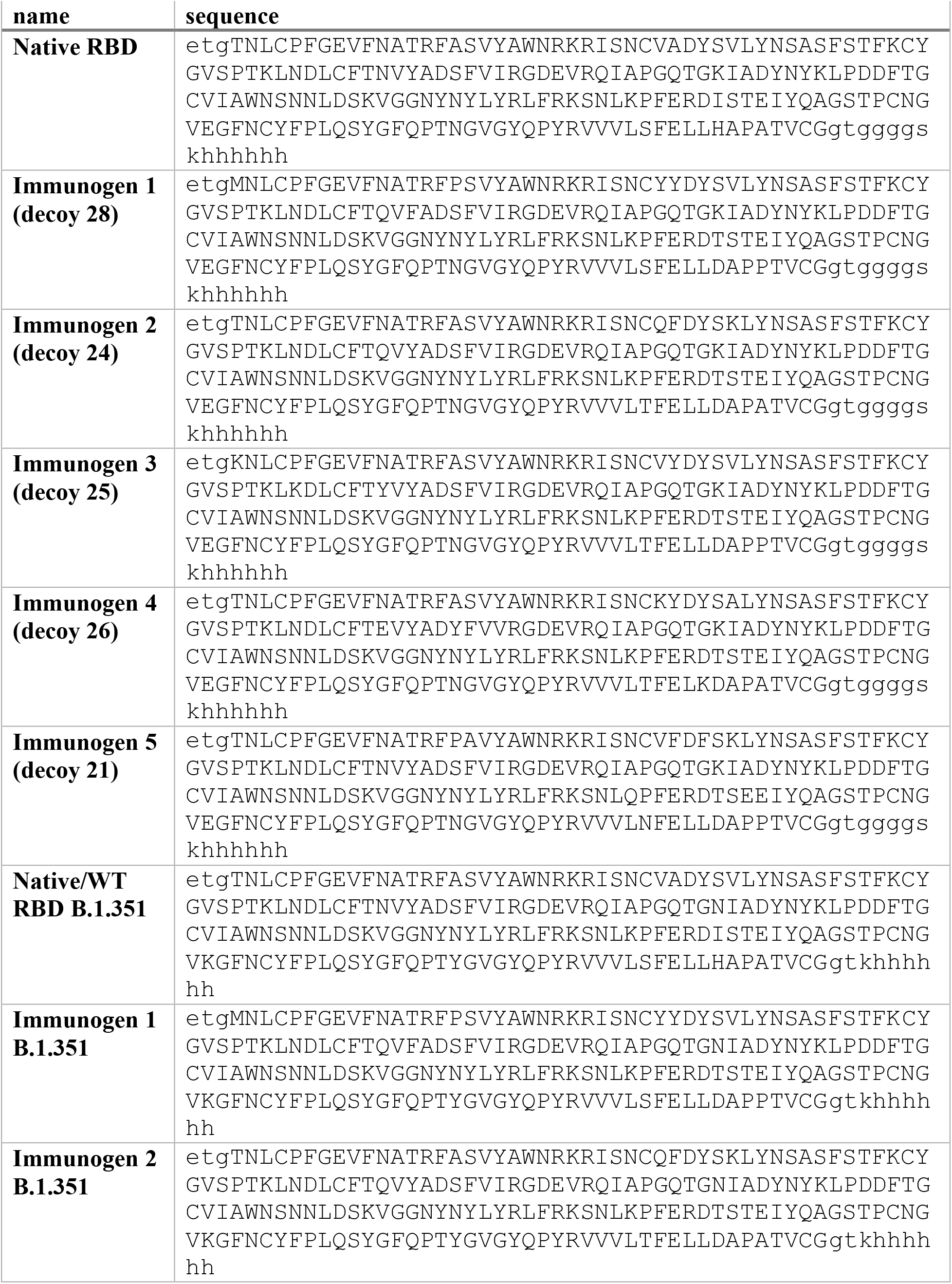
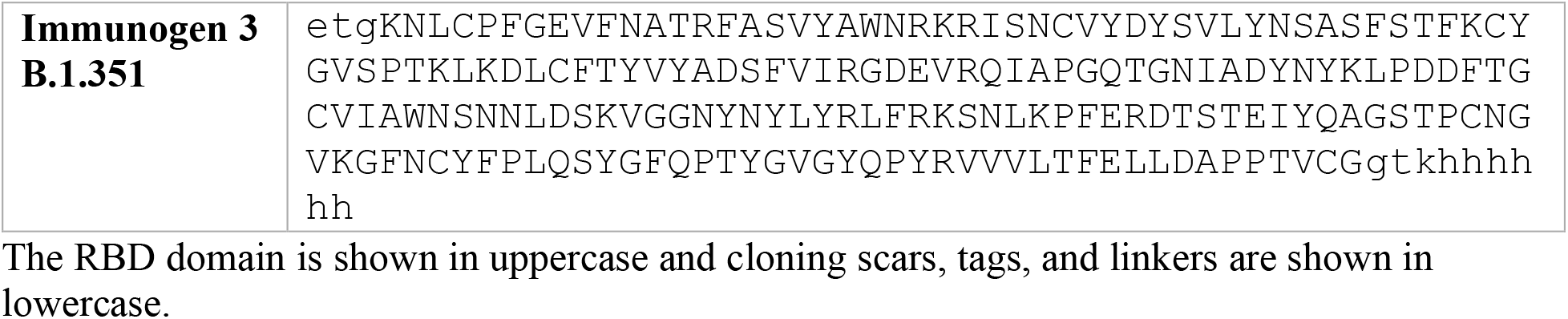
Sequences of lead immunogens after signal peptide cleavage.

**Extended Data Table 2.** Data collection and refinement statistics.

|  | Immunogen 3/P2B-2F6 |
| --- | --- |
| **Data collection** |  |
| Space group | P 21 21 21 |
| Cell dimensions |  |
| *a*, *b*, *c* (Å) | 71.06, 91.97, 111.15 |
| $\alpha$, $\beta$, $\gamma$ (°) | 90, 90, 90 |
| Resolution (Å) | 70.9 - 2.57 (2.66 - 2.57) * |
| *R*<sub>sym</sub> or *R*<sub>merge</sub> | 0.187 (2.089) |
| *I* / $\sigma I$ | 8.2 (1.02) |
| Completeness (%) | 99.9 (99.9) |
| Redundancy | 6.6 (6.8) |
| **Refinement** |  |
| Resolution (Å) | 56.23 - 2.6 (2.67 - 2.60) |
| No. reflections | 23032 (1472) |
| *R*<sub>work</sub> / *R*<sub>free</sub> | 0.221/0.270 (0.348/0.429) |
| No. atoms | 4861 |
| Protein | 4816 |
| Ligand/ion | 14 |
| Water | 31 |
| *B*-factors | 63.6 |
| Protein | 63.6 |
| Ligand/ion | 96.5 |
| Water | 49.9 |
| R.m.s. deviations |  |
| Bond lengths (Å) | 0.007 |
| Bond angles (°) | 1.12 |

One crystal was used for this dataset.
\*Values in parentheses are for highest-resolution shell.

**Extended Data Table 3.** Binding affinities of Ace2 and REGN10933 to RBD immunogens as determined by BLI.

|  | **K<sub>D</sub>**<br>(x 10<sup>-9</sup> ± SEM M) | **k<sub>a</sub>**<br>(x 10<sup>5</sup> ± SEM 1/Ms) | **k<sub>dis</sub>**<br>(x 10<sup>-3</sup> ± SEM 1/s) | **N** |
| --- | --- | --- | --- | --- |
| **Ace2 with WT-RBD** | 8.7 ± 0.9 | 7.7 ± 0.4 | 6.5 ± 0.4 | 3 |
| **Ace2 with immunogen 1** | 9.3 ± 0.7 | 8.0 ± 0.2 | 7.4 ± 0.4 | 3 |
| **Ace2 with immunogen 2** | 15 ± 3 | 7 ± 1 | 9.1 ± 0.4 | 3 |
| **Ace2 with immunogen 3** | 11 ± 4 | 8 ± 1 | 7.5 ± 0.8 | 3 |
| **Ace2 with immunogen 4** | 14 ± 3 | 6.6 ± 0.6 | 7.1 ± 0.9 | 3 |
| **REGN10933 with WT-RBD** | 1.8 ± 0.2 | 10.9 ± 0.8 | 1.9 ± 0.1 | 3 |
| **REGN10933 with immunogen 1** | 1.6 ± 0.1 | 12.2 ± 0.2 | 1.9 ± 0.1 | 3 |
| **REGN10933 with immunogen 2** | 1.4 ± 0.1 | 13.0 ± 0.3 | 1.8 ± 0.1 | 3 |
| **REGN10933 with immunogen 3** | 1.6 ± 0.1 | 11.7 ± 0.3 | 1.9 ± 0.1 | 3 |
| **REGN10933 with immunogen 4** | 1.5 ± 0.1 | 12.6 ± 0.2 | 1.8 ± 0.1 | 3 |

Binding data were fit using a 1:1 binding model. The averages and standard deviations of three biological replicates using independently purified immunogens are shown.

